# One-Entry Golden Seamless Cloning with a Reusable Donor Streamlines Plant Gene Functional Analysis

**DOI:** 10.64898/2026.10.05.752523

**Authors:** Yan Cheng, Yi Wu, Xiaona Ma, Kang Li, Luis Alejandro José Mur, Chuanying Fang

## Abstract

Molecular cloning underpins functional genomics, yet current toolboxes force a trade-off. Golden Gate frameworks handle one-pot multi-fragment assembly but overcomplicate routine single-gene cloning, whereas Gibson, In-Fusion, and Gateway systems need backbone linearization, resequencing, or proprietary recombinases that leave scars. Here we present Golden Seamless (GS), a one-entry, many-destination system in which a single PCR captures a gene once into a reusable Entry-and-Expression Clone that is never re-cloned. Circle-to-circle Golden Gate moves the same validated clone into plant, yeast, or bacterial vectors without linearization, and a *ccdB* counter-selection with orthogonal antibiotic markers drives background near zero. Bacterial destinations embed superfolder GFP, enabling naked-eye monitoring of recombinant proteins and an antibody-free dual-modality EMSA that resolves protein and DNA channels across a >150-nm spectral gap. GS thus consolidates capture, distribution, and validation into one low-background workflow with built-in quality control, cutting the time and cost of screening many candidates across functional assays.

## Dear Editor,

Molecular cloning is indispensable to functional genomics (Lee et al., 2008; Xing et al., 2016) and many cloning toolboxes have been developed, broadly categorized into direct-to-destination systems and entry-clone systems. Golden Gate frameworks (GreenGate, GoldenBraid, MoClo, and Mobius) excel at one-pot multi-fragment assembly but over-complicate routine single-gene cloning (Sarrion-Perdigones et al., 2011; Weber et al., 2011; Lampropoulos et al., 2013; Sarrion-Perdigones et al., 2013; Andreou and Nakayama, 2018). Gibson-based (Han et al., 2022) and In-Fusion-based (Luo et al., 2026) toolboxes use standardized homologous arms to build protein-protein interaction and localization constructs from a single primer pair, but they depend on linearized destination backbones and require resequencing of every final construct. Entry-clone systems are attractive because the insert is cloned once into a Gateway entry vector, which can shuffle the insert into multiple destination constructs without re-amplification and with a verified insert (Karimi et al., 2002; Luo et al., 2018). However, they depend on proprietary recombinases, leave long recombination-site scars, and produce an entry clone of no standalone utility. Here, we present Golden Seamless (GS), in which a single PCR captures the gene once into a reusable expressible vector (Entry and Expression Clone, EEC) that is never re-cloned. Every new assay simply feeds the same EEC into a plant, yeast, or bacterial vector through circle-to-circle Golden Gate, eliminating re-sequencing. Bacterial expression vectors incorporate superfolder GFP (sfGFP) fusions for direct monitoring of recombinant proteins and enable the development of dual-modality electrophoretic mobility shift assays (dmEMSA). This antibody-free method simultaneously visualizes proteins and DNA probes on a single native gel, allowing the direct detection of DNA-protein complexes.

We built three pGW7-backbone Golden Donor Vectors (GDVs; namely pY004, pY006 and pY009) carrying gentamicin and hygromycin resistance, an N-terminal 4×myc tag driven by a constitutive promoter, and a *Bsa*I-flanked *ccdB*–chloramphenicol module at the insertion site (Supplementary Methods). pY004 and pY006 both harbor the maize *ZmUBI* promoter, whereas pY009 carries the cauliflower mosaic virus (CaMV) *35S* promoter. In pY006 and pY009, inverted *Paq*CI sites adjacent to the *Bsa*I sites yield the same sticky ends. A single round of PCR appends homology arms or *Paq*CI sites to the gene of interest for insertion into GDVs via seamless or Golden Gate assembly, respectively; the resulting EEC is directly expressible in plants and, unlike the classical Gateway intermediate, retains both *Bsa*I sites flanking the insert (Fig. 1a and Table S1).

**Figure 1.**
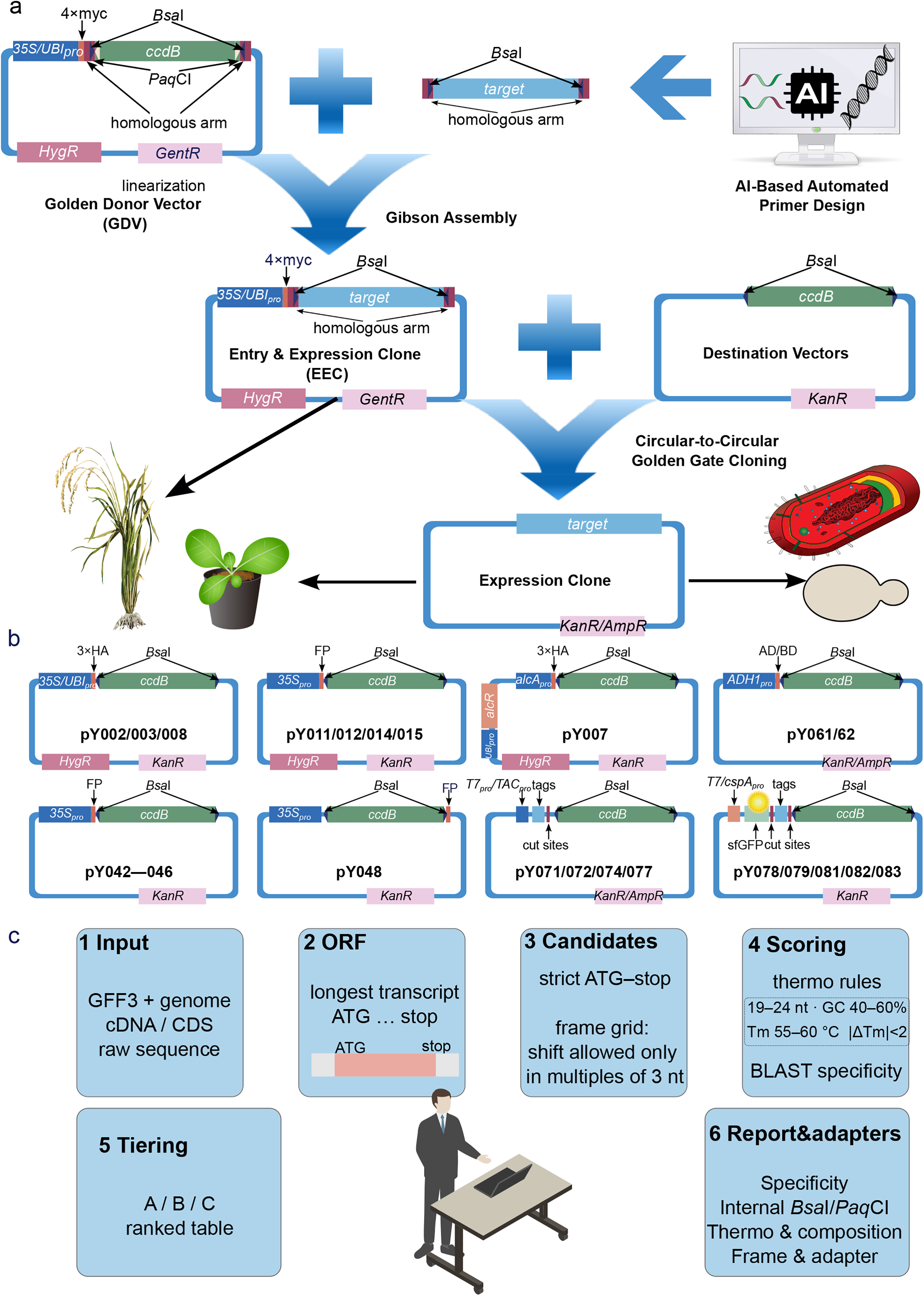
The Golden Seamless (GS) system: one-entry, many-destination cloning. **a)** Conceptual workflow. GSprimer designs primer pairs for amplification of a target gene. Each Golden Donor Vector (GDV) carries gentamicin and hygromycin resistance markers, an N-terminal 4×myc tag under a constitutive promoter, and a *Bsa*I-flanked *ccdB*–chloramphenicol module at the insertion site. Three GDVs were developed in this study: pY004 and pY006 both harbor the maize ubiquitin promoter (*UBI*_*pro*_), whereas pY009 carries the cauliflower mosaic virus (*CaMV*) 35S promoter (*35S*_*pro*_); pY006 and pY009 carry *Paq*CI sites in inverted orientation adjacent to the *Bsa*I sites, such that *Paq*CI digestion yields the same sticky ends as *Bsa*I, allowing both seamless and Golden Gate assembly, whereas pY004 supports seamless cloning only. A single round of PCR appends homology arms or *Paq*CI sites to the gene of interest for insertion into GDVs via seamless or Golden Gate assembly, respectively. The resulting Entry-and-Expression Clone (EEC) is directly expressible in plants and retains *Bsa*I sites flanking the insert. A one-pot, circle-to-circle Golden Gate reaction between the EEC and any circular destination vector excises the insert and ligates it into the destination backbone without linearization, producing an expression clone. Distinct antibiotic markers on the GDV and destination vectors, together with the *ccdB* counter-selection gene, drive the cloning background to near zero. **b)** Destination vector family. pCAMBIA1301-backbone vectors: pY002 (*35S*_*pro*_*:3×HA-GS*), pY003 (*UBI*_*pro*_*:3×HA-GS*), pY007 (*alcA*_*pro*_*:3×HA-GS*, ethanol-inducible), pY008 (*UBI*_*pro*_*:GS*, no tag), pY011 (*35S*_*pro*_*:mNeonGreen-GS*), pY012 (*35S*_*pro*_*:mScarlet-GS*), pY014 (*35S*_*pro*_*:nYFP-GS*), and pY015 (*35S*_*pro*_*:cYFP-GS*). pGreenII-backbone vectors: pY042 (*35S*_*pro*_*:nYFP-GS*), pY043 (*35S*_*pro*_*:cYFP-GS*), pY044 (*35S*_*pro*_*:mTurquoise2-GS*), pY045 (*35S*_*pro*_*:mNeonGreen-GS*), pY046 (*35S*_*pro*_*:mCherry-GS*), and pY048 (*35S*_*pro*_*:GS-mNeonGreen*). Yeast two-hybrid (Y2H) vectors: pY061 (pGADT7_GS) and pY062 (pGBKT7_GS). Bacterial expression vectors with the *T7* promoter (T7_*pro*_): pY071 (pET28a(+)_GS), pY072 (pET30a_GS), pY074 (pET49b_GS), pY077 (pGEX-6P-1_GS), pY078 (pET28-GG_GS, *N-sfGFP* incorporated), and pY079 (pET28-GH_GS, *N-sfGFP* incorporated). Bacterial expression vectors with the *cspA* promoter (*cspA*_*pro*_) and *N-sfGFP*: pY081 (*cspA*_*pro*_:*sfGFP*-*TEV-GST-HRV3C-GS*), pY082 (*cspA*_*pro*_:*sfGFP*-*TEV-His-HRV3C-GS*), and pY083 (*cspA*_*pro*_:*sfGFP*-*TEV-S tag-HRV3C-GS*). TEV (tobacco etch virus), SUMO (small ubiquitin-like modifier), HRV 3C (human rhinovirus 3C), or thrombin cleavage sites were integrated into the bacterial expression vectors, enabling precise tag excision. FP, fluorescent protein. **c)** GSprimer workflow. Given a gene identifier and a local genome annotation, GSprimer designs CDS primers anchored at the ATG/stop codon with GS adapter overhangs, verifies in-frame fusion with the 4×myc tag, screens primer specificity genome-wide, grades primer pairs into Tiers A/B/C, scans for internal *Bsa*I sites with synonymous wobble rescue, and recommends PCR conditions, making the entry PCR the only manual design step.

All GS destination vectors were generated by recombining a common GS element (a DNA block carrying the universal adapters, the two *Bsa*I sites, and *ccdB*) onto existing backbones. In a single Golden Gate assembly, the insert is excised from the sequencing-validated circular EEC and simultaneously ligated into any circular destination vector containing the same adapter sequence, obviating vector linearization. Distinct antibiotic resistance markers in the GDV and destination vectors, together with a *ccdB* counter-selection cassette lethal to standard cloning strains, minimize background carryover. Once captured and sequence-validated in an EEC, a gene can be rapidly mobilized into new expression, localization, or interaction constructs without re-cloning or re-sequencing.

We built pCAMBIA1301-derived vectors for stable overexpression with N-terminal tags (untagged or 3×HA) and fluorescent fusions (mNeonGreen, mScarlet, nYFP, cYFP), with an ethanol-inducible vector (pY007) for on/off comparisons in the same transgenic line. We modified pGreenII-62sk for protoplast transformation to carry fluorescent fusions (N-nYFP/cYFP/mTurquoise2/mNeonGreen/mCherry, C-mNeonGreen) and generated yeast two-hybrid (Y2H) and bacterial expression vectors by adding the GS element to pGADT7, pGBKT7, pET28a, pET30a, pET49b, and pGEX-6P-1 (Fig. 1b and Table S1). GSprimer automates primer design based on a given gene identifier and local genome annotation. The software anchors primers at the start and stop codons, verifies an in-frame fusion using the 4×myc tag, screens genome-wide specificity, and rescues internal *Bsa*I sites by synonymous wobble (Supplemental Video S1). This leaves the entry PCR as the only manual step (Fig. 1c and Table S1).

We validated the workflow and vectors for subcellular localization using rice calmodulin *OsCaM61*, in which an N-terminal GFP tag confers membrane association whereas a C-terminal GFP tag directs nucleoplasmic localization (Dong et al., 2002) (Fig. S1). For reliable subcellular colocalization, we characterized a rice bidirectional promoter (Fig. S2) that drives both fusions from one regulatory unit. Bidirectional expression was confirmed in *N. benthamiana*, and adding the minimal 35S promoter (*mini35S*) further enhanced and equalized the paired signals (Fig. S3). *mTurquoise2* inserted into pY007 showed a rapid and dose-dependent ethanol response (Fig. S4). We also validated the bimolecular fluorescence complementation (BiFC; Fig. S5) vectors with the homodimeric OsbZIP83 (Kobayashi et al., 2022) and bacterial expression vectors with the G-box-binding transcription factor OsMYL1 (Li et al., 2021) (Fig. S6).

Leveraging the robust folding, rapid maturation, bright fluorescence, and compatibility of sfGFP with N-terminal fusion (Pédelacq et al., 2006), we incorporated sfGFP into the bacterial destination vectors. The fusion protein emitted naked-eye-visible green fluorescence in culture, pellet, extract, and purified product (Fig. 2a–2d), enabling immediate visual identification of successful assemblies and expression. This feature facilitates the rapid exclusion of failed constructs and streamlines downstream expression optimization.

**Figure 2.**
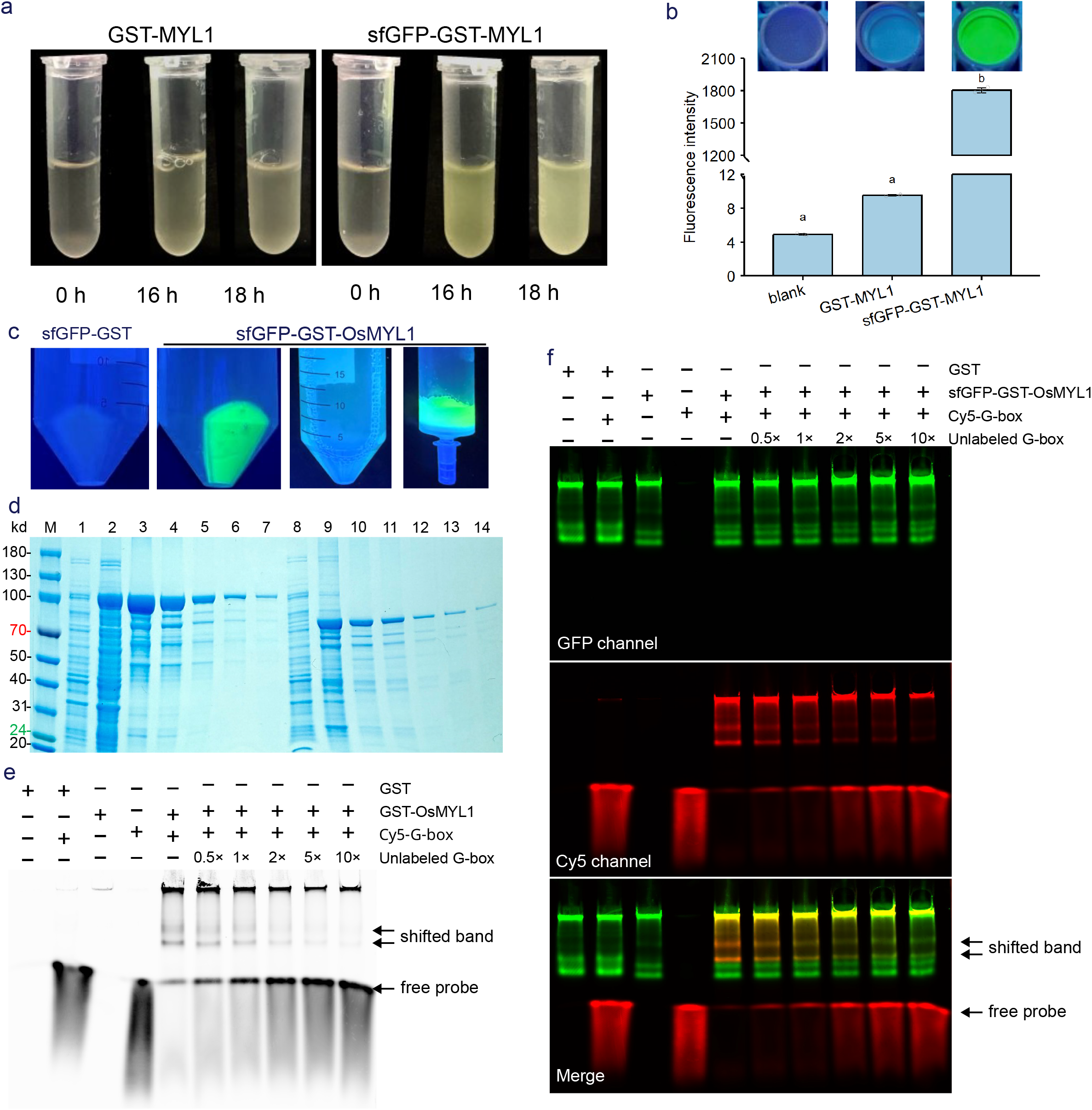
The sfGFP-tagged bacterial branch of GS enables naked-eye monitoring and purification tracking of recombinant OsMYL1 and underpins a dual-modality EMSA (dmEMSA). **a)** Culture phenotypes of GST–OsMYL1 and sfGFP–GST–OsMYL1 at 0, 16, and 18 h after induction. Only the sfGFP fusion emits visible green fluorescence following induction, whereas the GST–OsMYL1 culture remains non-fluorescent throughout. **b)** Fluorescence intensity of buffer (blank), GST–OsMYL1, and sfGFP–GST–OsMYL1 cultures at 18 h post-induction (n = 3). OD_600_ of each culture was adjusted to 0.3. Bars denote means; error bars denote standard deviation (SD); *p* < 0.05 by one-way ANOVA. **c)** Naked-eye visualization under a handheld UV flashlight. From left to right: (i) pelleted GST–OsMYL1 cells, (ii) pelleted sfGFP–GST–OsMYL1 cells, (iii) the flow-through from sfGFP–GST–OsMYL1 purification, and (iv) the purified sfGFP–GST–OsMYL1 product retained in a gravity-flow column. **d**) SDS-PAGE analysis of the expression and purification of GST–OsMYL1 and sfGFP–GST–OsMYL1. Lanes 1–14: 1, uninduced sfGFP–GST–OsMYL1; 2, total protein of sfGFP–GST–OsMYL1; 3, supernatant of sfGFP–GST–OsMYL1; 4–7, purified sfGFP–GST–OsMYL1 fractions; 8, uninduced GST–OsMYL1; 9, total protein of GST–OsMYL1; 10, supernatant of GST–OsMYL1; 11–14, purified GST–OsMYL1 fractions. **e)** EMSA. A standard electrophoretic mobility shift assay confirms sequence-specific binding of GST–OsMYL1 to the G-box probe, yielding a shifted band that is competed off by abolished unlabeled probe. **f)** dmEMSA. The built-in sfGFP serves as an orthogonal protein channel while a Cy5-labeled G-box probe provides the DNA channel; the >150-nm spectral gap separates the green (protein) and red (probe) signals, allowing complex detection without antibodies, with the merged image showing co-migrating yellow at the shifted position.

Conventional green fluorescent EMSA probes emit in the same spectral region as sfGFP (∼509 nm), reventing clear discrimination between probe and fusion protein (Fig. S7). Thus, we repurposed sfGFP for dmEMSA by pairing it with a spectrally orthogonal fluorophore (here Cy5, ∼667 nm). Our dmEMSA exploits a >150-nm spectral gap to resolve the protein and DNA channels and detect binding without antibodies, which represents an improvement over other EMSA schemes (Demczuk et al., 1993; Sumner et al., 2026). Our strategy was validated on a native gel, in which sfGFP-fused OsMYL1 and the Cy5-labeled G-box probe appeared as green and red signals, respectively, and the shifted complex appeared yellow. This interaction was outcompeted by a 5-fold excess of unlabeled probe (Fig. 2e, 2f).

By consolidating capture, distribution, and validation into one low-background workflow with a built-in quality-control readout, GS reduces the time and cost of testing many candidates across functional assays. This consolidation targets the step that limits plant functional analysis and aids dissection of agronomically important gene families and stress-response pathways. Vectors become GS-compatible by incorporating the GS element, extending the platform to diverse demands. Although internal *Bsa*I/*Paq*CI sites within the target gene theoretically constrain the system, these can be identified by GSprimer. In practice, one or two internal *Bsa*I/*Paq*CI sites are not an absolute barrier, and prolonging the Golden Gate reaction improved the positive-clone rate under these conditions.

## Author contributions

Conceptualization: C.F. Methodology: C.F., Y.C., and Y.W. Investigation: Y.C., Y.W., X.M. and K.L. Visualization: Y.C., Y.W., X.M., K.L., and C.F. Funding acquisition: C.F. Project administration: C.F. Supervision: C.F. Writing – original draft: C.F., Y.C., Y.W., and L.M. Writing – review & editing: C.F. and L.M.

## Acknowledgements

We thank Prof. Lei Yao from Beijing Academy of Agriculture and Forestry Sciences for providing pBBB-GRA-Cre/loxP plasmid.

## Supplementary material

Supplementary Table S1. Golden destination vectors and destination vectors in Golden Seamless Cloning.

Supplementary Table S2. Primers used in this study.

Figure S1. Validation of the pGreenII transient backbone by OsCaM61 subcellular localization.

Figure S2. Identification of a rice bidirectional promoter *pBD*.

Figure S3. Functional validation of the bidirectional promoter (*pBD*) in living cells.

Figure S4. Transient-expression validation of the ethanol-inducible overexpression vector in *N. benthamiana*.

Figure S5. Validation of the protein–protein interaction destination vectors with OsbZIP83 self-interaction.

Figure S6. EMSA with non-sfGFP OsMYL1.

Figure S7. EMSA with sfGFP-OsMYL1 and FAM-modified probe.

Supplemental Video S1. Step-by-step workflow of CDS primer design with GSprimer, demonstrated using the rice gene *OsbZIP83* (*LOC_Os12g05680*).

Supplementary Dataset S1. The fasta file of the bidirectional promoter (pBD.fasta).

## Data availability

All data generated or analyzed during this study are included in this published article and its sasupplementary information files. The GSprimer source code and documentation are freely available at https://github.com/ChuanyingFang/GSprimer. The derived vectors and associated protocols are available for non-commercial purposes upon request from the corresponding author.

