## Supplementary Tables,Supplementary Figures,Supplementary Methods, and Supplementary Video for "One-Entry Golden Seamless Cloning with a Reusable Donor Streamlines Plant Gene Functional Analysis": Supplementary Tables-biorxv.pdf

**Supplementary Table S1**. Golden destination vectors and destination vectors in Golden Seamless Cloning.

| Name | Construct | Purpose | Marker | Purpose |
|---|---|---|---|---|
| pY004 | pY004-pGW7_GSHUMC_G | GDV | gent | UBI$_{pro}$, N-4×myc |
| pY006 | pY006-pGW7_GSHUMC_G | GDV | gent | UBI$_{pro}$, N-4×myc |
| pY009 | pY009-pGW7_GSHCMC_G | GDV | gent | 35S$_{pro}$, N-4×myc |
| pY002 | pY002-pCAMBIA1301_GSHCHC_K | OE_cons | Kn | 35S$_{pro}$, N-3×HA |
| pY003 | pY003-pCAMBIA1301_GSHUHC_K | OE_cons | Kn | UBI$_{pro}$, N-3×HA |
| pY007 | pY007-pCAMBIA1301_GSHAHC_K | OE_indu | Kn | N-3×HA, ethanol-inducible |
| pY008 | pY008-pCAMBIA1301_GSHUC_K | OE_cons | Kn | UBI$_{pro}$, no tags |
| pY011 | pY011-pCAMBIA1301_GSHCmNGC_K | subcellular | Kn | 35S$_{pro}$, N-mNeonGreen |
| pY012 | pY012-pCAMBIA1301_GSHCmSC_K | subcellular | Kn | 35S$_{pro}$, N-mScarlet |
| pY014 | pY014-pCAMBIA1301_GSHCnYC_K | BiFC | Kn | 35S$_{pro}$, N-nYFP |
| pY015 | pY015-pCAMBIA1301_GSHCcYC_K | BiFC | Kn | 35S$_{pro}$, N-cYFP |
| pY042 | pY042-pGreen II 62sk_GSTCnYC_K | BiFC | Kn | 35S$_{pro}$, N-nYFP |
| pY043 | pY043-pGreen II 62sk_GSTCcYC_K | BiFC | Kn | 35S$_{pro}$, N-cYFP |
| pY044 | pY044-pGreen II 62sk_GSTCTC_K | subcellular | Kn | 35S$_{pro}$, N-mTurquoise2 |
| pY045 | pY045-pGreen II 62sk_GSTCGC_K | subcellular | Kn | 35S$_{pro}$, N-mNeonGreen |
| pY046 | pY046-pGreen II 62sk_GSTCCC_K | subcellular | Kn | 35S$_{pro}$, N-mCherry |
| pY048 | pY048-pGreen II 62sk_GSTCCG_K | subcellular | Kn | 35S$_{pro}$, C-mNeonGreen |
| pY061 | pY061-pGADT7_GSAAC_A | YH | Amp | Yeast one/two hybrid |
| pY062 | pY062-pGBKT7_GSABC_K | YH | Kn | Yeast two hybrid |
| pY071 | pY071-pET28A(+)_GS_K | Prot Expr | Kn | N-His, Thrombin, T7; C-His |
| pY072 | pY072-pET30a_GS_K | Prot Expr | Kn | N-His, S, Thrombin, EK; C-His |
| pY074 | pY074-pET49b_GS_K | Prot Expr | Kn | N-GST, His, HRV 3C; C-S, Thrombin |
| pY077 | pY077-pGEX-6P-1_GS_A | Prot Expr | Amp | N-GST, HRV3C |
| pY078 | pY078-pET28-GG_GS_K | Prot Expr | Kn | N-His, Thrombin, T7, N-His, Thrombin, T7, sfGFP, TEV, GST, HRV3C; C-His |
| pY079 | pY079-pET28-GH_GS_K | Prot Expr | Kn | N-His, Thrombin, T7, sfGFP, TEV, His, HRV3C; His |
| pY081 | pY081-pCold-GG_GS_K | Prot Expr | Kn | N-sfGFP, TEV, GST, HRV3C |
| pY082 | pY082-pCold-GH_GS_K | Prot Expr | Kn | N-sfGFP, TEV, His, HRV3C |
| pY083 | pY083-pCold-GS_GS_K | Prot Expr | Kn | N-sfGFP, TEV, S, HRV3C |

GDV: Golden Donor Vector; OE_cons: Constitutive overexpression; OE_indu: Induced overexpression; Prot Expr: Protein Expression.

**Supplementary Table S2**. Primers used in this study.

| Name | Sequence (5'-3') |
|---|---|
| GM-F | cacgttaagggattttgttaggtggcggtacttggg |
| GM-R | atgttatggagcagcaacgatgttacgcagcagcaacg |
| UBI-F | cgttgctgctccataacatcaa |
| UBI-R | ctcgagagacctgatatcactagtcagatcctcttcagagat |
| BKA-F | tcgttgctgctccataacatcaaacatcgacccacggc |
| BKA-R | cgggtcacgctgcactgcaggcatgcgagctcgaattatcatacatgag |
| BKB-F | ccaaaatcccttaacgtgagttttcgttcca |
| BKB-R | ggattgagaccacccgcggccatgctagagtccgcaaaaat |
| 4GS-F | gtgatatcaggtctctcgaggtaggcaggtgacttagtcgaccgacagccttc |
| 4GS-R | gccgcgggtggtctcaatccgtaggcaggtgcatcacgacgcactttgc |
| 35-F | gtatgataattcgagctcggagaggcggtttgcgtattgg |
| 35-R | agatgagtttctgctccatatctcattgccccccggatctgcga |
| 4m-F | tcgcagatccggggggcaatgagatatggagcagaaactcatct |
| 4m-R | ctcgagagacctgatatcactagtcagatcctcttcagagatgag |
| LC-F | gtgatatcaggtctctcgagaataaatacctgtgacggaag |
| LC-R | gccgcgggtggtctcaatccgcctactcgctattgtcctca |
| PJG066-F | aggtcgaaaaggtttctttcctgtggatagcacgtacatt |
| PJG066-R | aatgtacgtgctatccacaggaaagaaaccttttcgacct |
| BKC-F | acgttgtaaaacgacggccagtgccaagcttg |
| BKC-R | aggatccgagctcgaatttccccgatcgttcaaacatttggcaataaag |
| GSC-F | cgttccagattacgctggtaccctcgagagagaccaataaatacctgtgac |
| GSC-R | aaattcgagctcggatcctgagaccgcctactcgctattgtcctc |
| SHR1-F | gtaaaacgacggccagtgccaagcttggagaggcggtttgcgtattggc |
| SHR1-R | acgtcatacggatagcccgcatagtcaggaacatcgtatgggtacatactagtatctcattgccccccgga |
| SHR2-F | gtaaaacgacggccagtgccaagcttggagaggcggtttgcgtattggc |
| SHR2-R | ctcgagggtaccagcgtaatctggaacgtcatatggataggaccctgcatagtccgggacgtcatacggatagccc |
| 2NG-F | ggggcaatgagatactagtatgacttcgaagggtgaagaag |
| 2NG-R | tggtctctctcgagggtacccttataaagctcatccattccc |
| 2mS-F | ggggcaatgagatactagtatggtgagcaagggcgaggcag |
| 2mS-R | attggtctctctcgagggtacccttgtacagctcgtccatgc |
| 2nY-F | gggggcaatgagatactagtatggtgagcaagggcgagga |
| 2nY-R | ttggtctctctcgagggtaccgtcctcgatgttgtggcgga |
| 2cY-F | gggggcaatgagatactagtatgggcggcagcgtgcagct |
| 2cY-R | tggtctctctcgagggtacccttgtacagctcgtccatgc |
| EIO-F | aggtcgactctagactagtatggcagatacgcgccgacg |
| EIO-R | gtatgggtacatactagtgtcctctccaaatgaaatgaac |
| 62NGS-F | aacacctgctgaacgagagagaccaataaatacct |
| 62NGS-R | aacacctgctgaaagcttatcctgagaccgcctactcgc |

| | |
|---|---|
| 62CGS-F | gacacctgcgtcgctagtcgagagagaccaataaatacctg |
| 62CGS-R | gacacctgcgtcgggatcctgagaccgcctact |
| 62nY-F | aacacctgctgaactagtatggtgagcaagggcgag |
| 62nY-R | aacacctgctgaacgaggtcctcgatgttgtggcg |
| 62cY-F | aacacctgctgaactagtatgggcggcagcgtgcag |
| 62cY-R | aacacctgctgaacgagcttgtacagctcgtccatgccg |
| 62mT-F | gcggccgctctagaactagtatggtgagcaagggcgagg |
| 62mT-R | tggtctctctcgaggatatccttgtacagctcgtccatgccg |
| 62NG-F | gcggccgctctagaactagtatgacttcgaagggtgaagaag |
| 62NG-R | tggtctctctcgaggatatccttataaagctcatccattcccat |
| 62mC-F | gcggccgctctagaactagtatggtgagcaagggcgagg |
| 62mC-R | tggtctctctcgaggatatccttgtacagctcgtccatgccg |
| 62CNG-F | gacacctgcgtcgatccatgacttcgaagggtgaagaa |
| 62CNG-R | gacacctgcgtcgagctttcacttataaagctcatccattcccat |
| AGS-F | atggaggccagtgaattactcgagagagaccaataaatacct |
| AGS-R | gagctcgatggatcaggatcctgagaccgcctactcgct |
| BGS-F | catggaggccgaattagagctcgagagagaccaataaatacct |
| BGS-R | tgcaggtcgacggatcaggatcctgagaccgcctactcgct |
| 28GS-F | atccgaattcgagctcctcgagagagaccaataaatacc |
| 28GS-R | tggtggtggtgctcgagggatcctgagaccgcctactcgc |
| 30GS-F | aaggccatggctgatctcgagagagaccaataaatacc |
| 30GS-R | cgaattcggatccgatgatcctgagaccgcctactcgcta |
| 49GS-F | ttcagggacccgggtacctcgagagagaccaataaatacc |
| 49GS-R | cagaattcggatcggatcctgagaccgcctactcgctat |
| 6PGS-F | ggcccctgggatccccggaattcctcgagagagaccaataaatac |
| 6PGS-R | tcacgatgcggccgctcgatatgatcctgagaccgcctactcgctattgtcc |
| sf-F | tgaatcacaaagtgactagtatgagcaaaggcgaagaactg |
| sf-R | ggattggaagtacaggttttctccatagccatgagtgatacccgcc |
| CGG-F | gaaaacctgtacttccaatccaagctttcccctatactaggttattggaaaatta |
| CGG-R | tggcagggacttctctagaatcctgagaccgcctactcgc |
| HGS1-F | ctcgagctggaagttctgttccaggggcccctcgagtgagaccaataaatacc |
| HGS2-F | agcacctgcggcaagcttcatcaccatcaccatcacctcgagctggaagttctgttc |
| HGS-R | aggcacctgcggcactagaatcctgagaccgcctactcgc |
| SGS1-F | ctcgagctggaagttctgttccaggggcccctcgagtgagaccaataaatacc |
| SGS2-F | gctaaattcgaacgccagcacatggacagcctcgagctggaagttctgttc |
| SGS3-F | agcacctgcggcaagcttaaagaaaccgctgctgctaaattcgaacgccagcaca |
| SGS-R | aggcacctgcggcactagaatcctgagaccgcctactcgc |
| GTG-F | ccgaattcgagctcctcgagatgagcaaaggcgaagaactg |
| GTG-R | tggtggtggtggtgctcgaggaatcctgagaccgcctactc |
| PA-F | agggtctcagaagtgagacttttcaacaaagggt |

| | |
|---|---|
| PA-R | agggtctcaagcttgatctagtaacatagatgacac |
| PB-F | agggtctcactagtatgacttcgaagggtgaagaag |
| PB-R | agggtctcacttcccaacagttgcgcagc |
| PC-F | agggtctcactagtctctctagctgcaatagaaaac |
| PC-R | agggtctcactagtgcacgaaaggcacaaaaaaac |
| PD-F | agggtctcaaattcccgatctagtaacatagatg |
| PD-R | agggtctcactagtatggtgagcaagggcgaggagc |
| MBM-F | agagtcgacctgcactagtgtcctctccaaatgaaatgaacttccttatatagactctctagctgcaatagaaaac |
| MBM-R | cctcgcccttgctcaccatactagtgtcctctccaaatgaaatgaacttccttatatagagcacgaaaggcacaaaaaaac |
| OsCaM61-F | gtgatatcaggtctctcgagtttgctcgccttctcgaag |
| OsCaM61-R | gccgcgggtggtctcaatcctcgtgtagctcattgactca |
| OsCaM61-R2 | gccgcgggtggtctcaatcccaggatcacgcacttctgg |
| OsMYL1-F | gtgatatcaggtctctcgagatgtcgtggtccgagacg |
| OsMYL1-R | gccgcgggtggtctcaatcctcacggggagggagtgg |
| OsbZIP83-F | agcacctgcagtccgagactagtatgggagaggctaggagagg |
| OsbZIP83-R | agcacctgcagtcatcctcagaaagctgaaaattggctttc |
| G-box probe | atgtatttgtatttgagcacgtgttaaattagctcttac |
