## Supplementary Tables,Supplementary Figures,Supplementary Methods, and Supplementary Video for "One-Entry Golden Seamless Cloning with a Reusable Donor Streamlines Plant Gene Functional Analysis": SupplementaryFigures261003.pdf

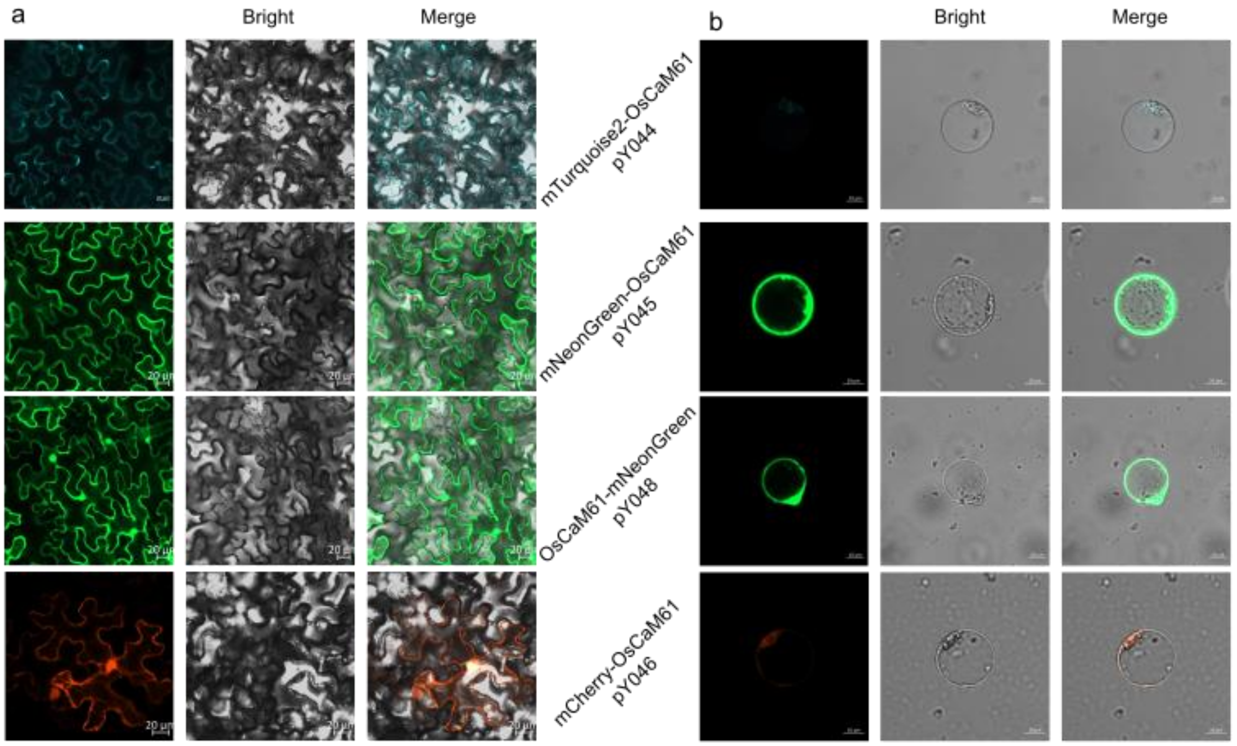


**Figure S1**. Validation of the pGreenII transient backbone by OsCaM61 subcellular localization. The *OsCaM61* CDS was cloned into the Golden Donor Vector pY004 and transferred by Golden Gate into the pGreenII-backbone transient expression destination vectors pY044 (N-mTurquoise2), pY045 (N-mNeonGreen), pY046 (N-mCherry), and pY048 (C-mNeonGreen). The expression clones were transiently expressed in a) *N. benthamiana* leaves and b) rice protoplasts. Confocal images were acquired on a Carl Zeiss LSM 980 microscope. Scale bars, 20 µm.

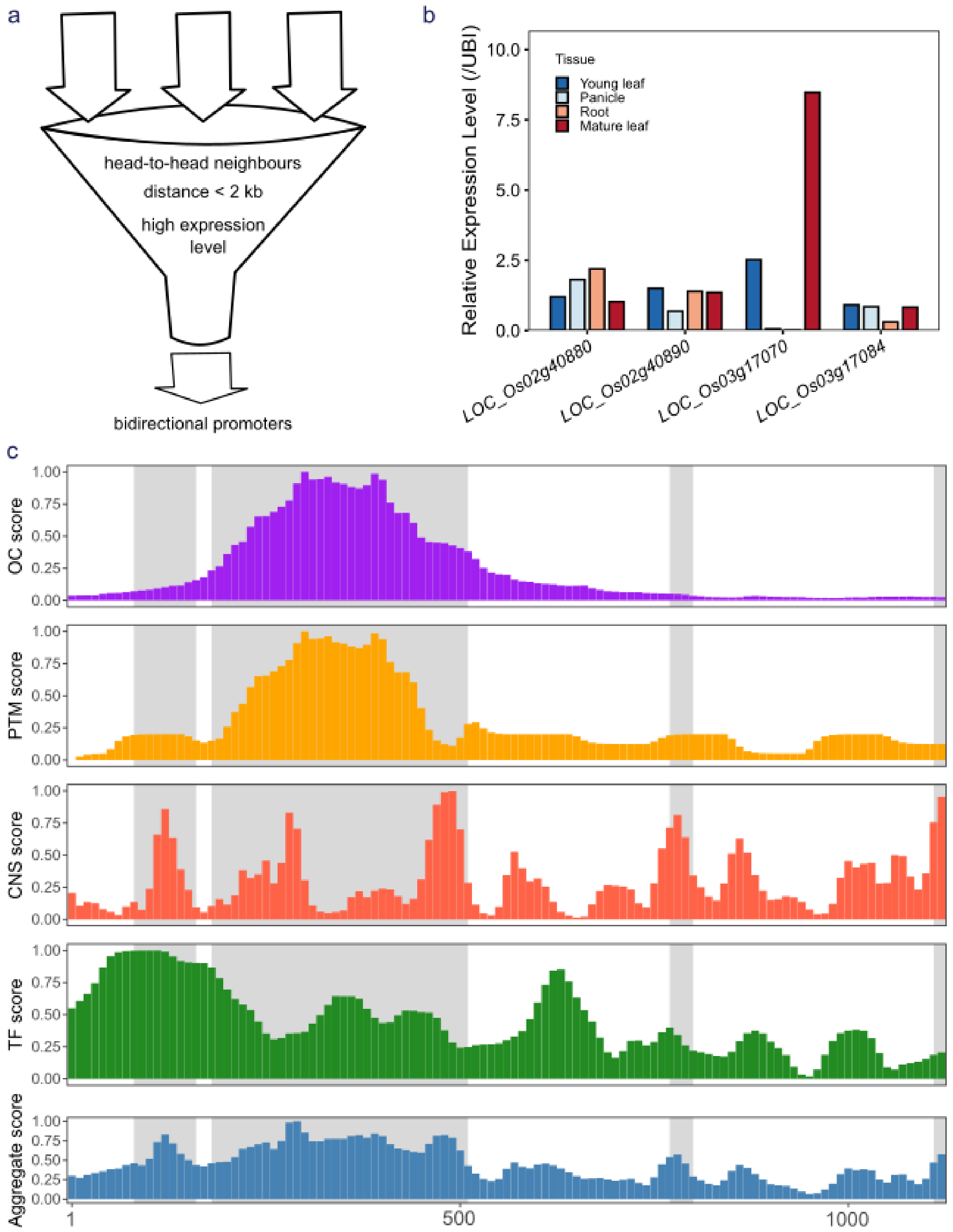


**Figure S2**. Identification of a rice bidirectional promoter *pBD*. a) Schematic diagram of the identification pipeline. Bidirectional promoters were mined from *RiceSuperPIRdb* based on three criteria: head-to-head adjacent gene pairs located on opposite strands, transcription start sites within a 2 kb interval, and expression levels comparable to those of the rice reference gene *OsUbq*

(*LOC_Os03g13170*). b) Candidate gene expression levels. Expression data were retrieved from *RiceSuperPIRdb*. c) Feature and aggregate scores of *pBD* calculated using the CAPE model. CAPE integrates multiple regulatory features, including open chromatin (OC), histone post-translational modifications (PTMs), conserved non-coding sequences (CNSs), and transcription factor binding sites (TFs), to comprehensively assess the editability of the target region. From top to bottom, the tracks represent the OC, PTM, CNS, and TF feature scores, followed by the aggregate score. The x-axis indicates positions within the sequence, and the y-axis represents the normalized score for each feature. Gray-shaded indicate key editable regions predicted by CAPE.

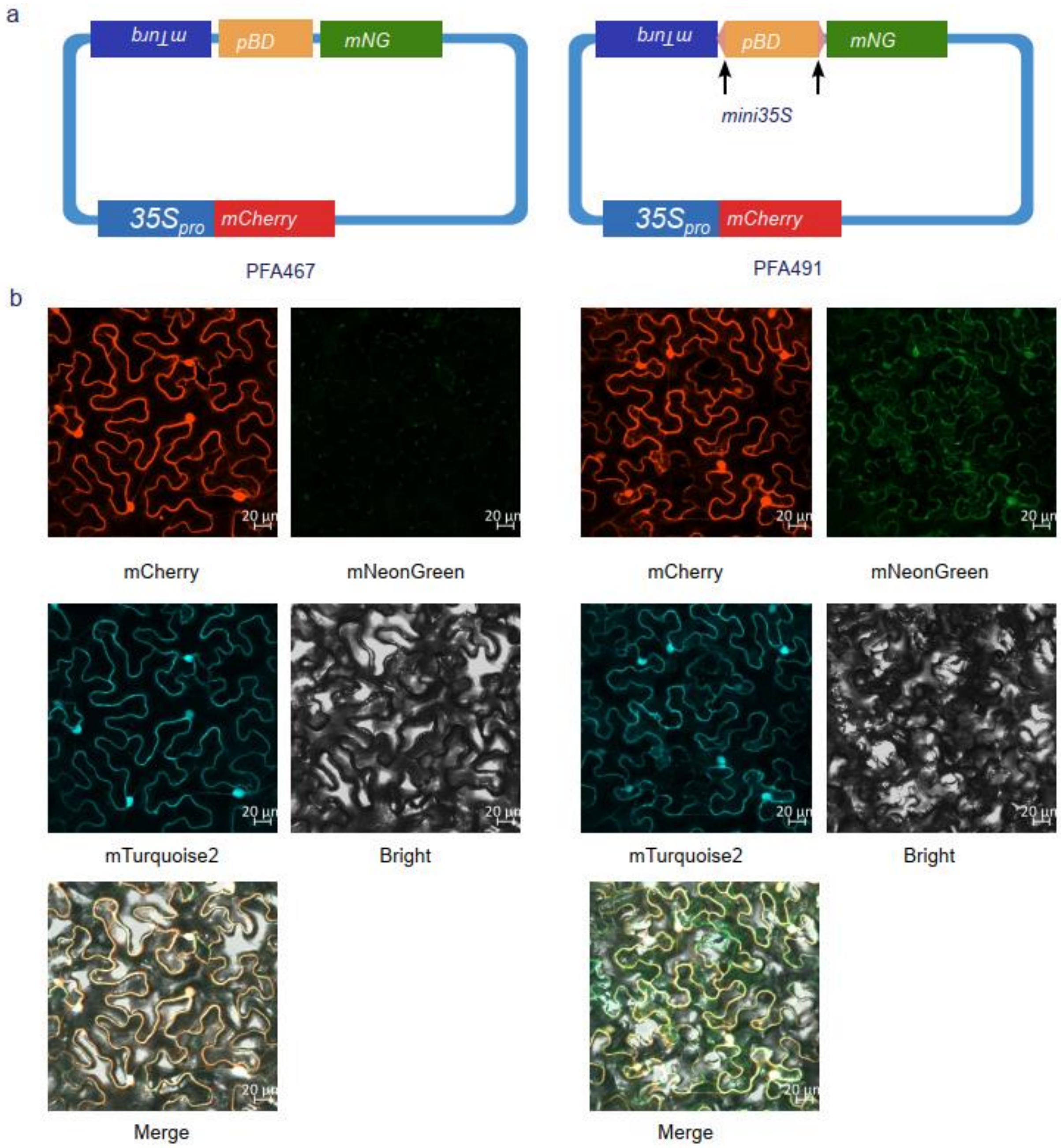


**Figure S3**. Functional validation of the bidirectional promoter (*pBD*) in living cells. a) Schematic of the dual fluorescent protein system driven by *pBD* (PFA467) and the bidirectional *mini35S*-fused *pBD* variant (PFA491). *mTurquoise2* and *mNeonGreen* are expressed in opposite directions by *pBD*, while the CaMV *35S* promoter (*35S*$_{pro}$) drives *mCherry* as an internal control. b) Fluorescence was visualized in *N. benthamiana* leaves after *Agrobacterium*-mediated infiltration using confocal laser scanning microscopy. Scale bars = 20 µm.

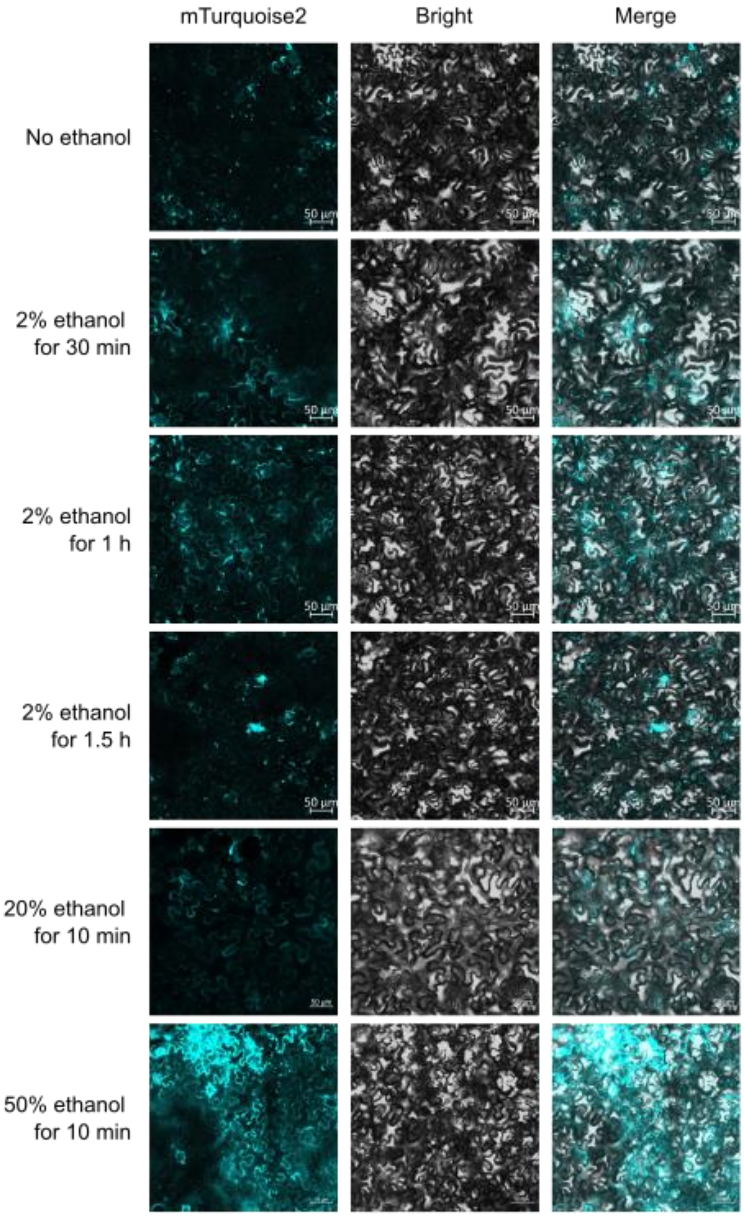


**Figure S4**. Transient-expression validation of the ethanol-inducible overexpression vector in *N. benthamiana*. *mTurquoise2* was inserted into the ethanol-inducible destination vector pY007 (*$UBI_{pro}$-alcR-$35S_{Ter}$-$alcA_{pro}$-mini35S*) and expressed under the control of *$alcA_{pro}$-mini35S*. *Agrobacterium tumefaciens* carrying the construct was infiltrated into *N. benthamiana* leaves. After 24 h of dark incubation followed by 16 h of light, four treatments were applied topically to the adaxial surface of living leaves: water (no ethanol, CK) or 2%, 20%, or 50% ethanol (50–100 µL each). Leaf discs (0.5

× 0.5 cm) were harvested at 30 min, 1 h, and 1.5 h after 2% ethanol treatment and 10 min after 20% or 50% ethanol treatment. Confocal images were acquired on a Carl Zeiss LSM 980 microscope. Scale bars, 50 μm.

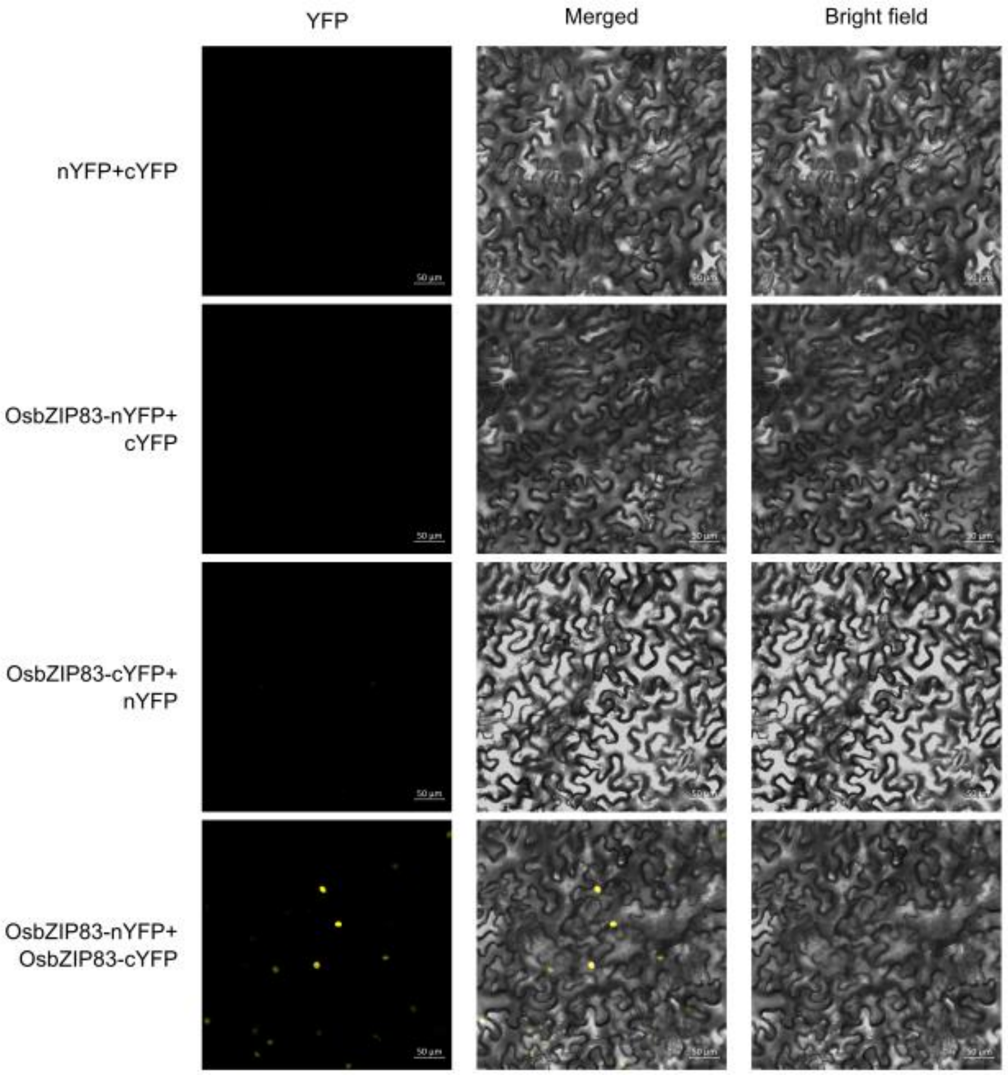


**Figure S5**. Validation of the protein–protein interaction destination vectors with OsbZIP83 self-interaction. The CDS of *OsbZIP83* was cloned into BiFC destination vectors based on pCAMBIA1301 backbones (pY014 and pY015), fused with nYFP and cYFP, respectively. The resulting expression clones were introduced into *N. benthamiana* leaves. Confocal images were acquired on a Carl Zeiss LSM 980 microscope. Scale bars, 50 μm.

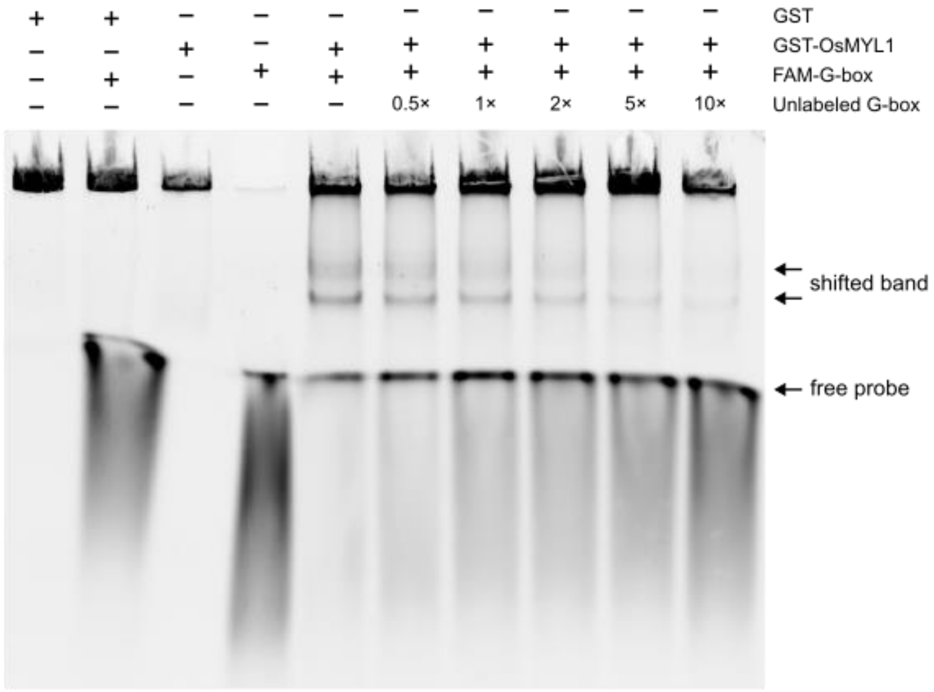


**Figure S6**. EMSA with non-sfGFP OsMYL1. EMSA was performed with GST–OsMYL1 and a FAM-modified G-box probe (the same probe used for dmEMSA). The shifted band was abolished by excess unlabeled probe, confirming sequence-specific binding. The gel was imaged directly on a GE Amersham Typhoon 5 scanner. A single channel was acquired with a 488-nm laser and the Cy2 525BP20 emission filter. Images were acquired in normal-scan mode (100 µm pixel size) to minimize background fluorescence.

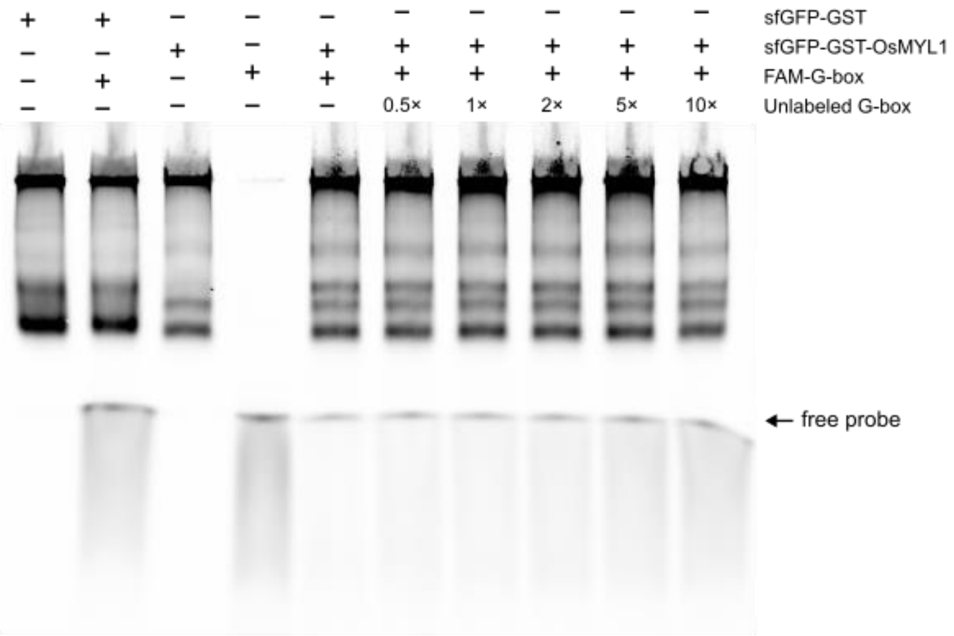


**Figure S7**. EMSA with sfGFP-OsMYL1 and FAM-modified probe. sfGFP–OsMYL1 was incubated with a FAM-modified G-box probe (the same probe as in Fig. S6). In a single-channel acquisition, FAM emission overlaps the sfGFP detection window, so the probe and protein signals cannot be resolved—the limitation that dmEMSA overcomes. The gel was imaged directly on a GE Amersham Typhoon 5 scanner. A single channel was acquired with a 488-nm laser and the Cy2 525BP20 emission filter. Images were acquired in normal-scan mode (100 µm pixel size) to minimize background fluorescence.
