## Supplementary Tables,Supplementary Figures,Supplementary Methods, and Supplementary Video for "One-Entry Golden Seamless Cloning with a Reusable Donor Streamlines Plant Gene Functional Analysis": SupplementaryMethods261003.pdf

### ***Plant and bacterial materials***

Rice (*Oryza sativa* L. cv. Zhonghua 11) was used for protoplast isolation. *Agrobacterium tumefaciens* strains GV3101 (Weidi, Shanghai, China, Cat. No. AC1001) and AGL1 (pSoup-p19) (Weidi, Shanghai, China, Cat. No. AC1023) were used for *N. benthamiana* transformation. *Escherichia coli* strains Top10 (routine cloning; *ccdB*-sensitive; Weidi, Shanghai, China, Cat. No. DL1010), DB3.1 (maintenance of *ccdB* vectors; Weidi, Shanghai, China, Cat. No. DL1040), and BL21(DE3) (protein expression, Weidi, Shanghai, China, Cat. No. EC1002) were used as appropriate.

### ***DNA assembly***

All plasmids generated in this study—including Golden Donor Vector (GDV) construction, Entry-and-Expression Clone (EEC) generation, destination-vector engineering, and routine subcloning—were assembled with one of the following seamless-cloning or Golden Gate workflows.

For seamless cloning, each vector backbone was linearized at a single restriction site, with homology arms flanking the cloning site matching the ends of the donor PCR product. For the Gibson-type route, the linearized backbone was re-circularized with the donor insert in a one-step isothermal reaction (Uniclone One Step Seamless Cloning Kit, Genesand, Beijing, China, Cat. No. SC612; 15–25-nt homology arms; 50 °C for 60 min). For the In-Fusion-type route, the linearized backbone was recombined with the donor in a single-recombinase, ligase-free reaction (SE Seamless Cloning Kit, ZC231; SE Recombinase; 15–20-bp end homology; 37 °C for 60 min). In both routes, the donor insert was added at a 3-fold molar excess over 30 ng of linearized backbone; the mass of each insert was calculated as 30 ng × 3 × (insert length / backbone length), equivalent to fmol = ng × $10^6$ / (bp × 650).

Golden Gate assemblies were performed as single-tube restriction–ligation reactions using *Bsa*I-HF v2 or *Paq*CI type IIS enzymes with T4 DNA ligase. Each 5 µL reaction contained 50 ng of destination vector, the donor plasmid at a donor-to-destination molar ratio of 3:1 (calculated as above), 0.5 μL of 10× T4 DNA ligase buffer (NEB, Ipswich, MA, USA, Cat. No. B0202S), 0.1 μL of *Bsa*I-HF v2 (NEB, Ipswich, MA, USA, Cat. No. R3733V) or *Paq*CI (NEB, Ipswich, MA, USA, Cat. No. R0745SVIAL) supplemented with equal *Paq*CI activator (NEB, Ipswich, MA, USA, Cat. No. S0532SVIAL), 0.2 μL of T4 DNA ligase (NEB, Ipswich, MA, USA, Cat. No. M0202S), and nuclease-free water to 5 μL. Reactions were incubated at 37 °C for 30 s first, then cycled 35 times between 37 °C for 3 minutes and 16 °C for 4 minutes, followed by 50 °C for 5 min, 80 °C for 10 min, and a hold at 4 °C.

For multi-insert assemblies, all inserts were included at equimolar ratios, with individual masses scaled by fragment length to maintain stoichiometry. Donor ratios assume a single excised insert per donor plasmid; for donors carrying multiple *Bsa*I-flanked modules, molar amounts were calculated for each excised fragment.

Reaction products were transformed into chemically competent *E. coli* DB3.1 or Top10 (Weidi, Shanghai, China); constructs carrying the *ccdB* counter-selection cassette were propagated in DB3.1, whereas all other plasmids were maintained in Top10. Positive clones were screened by colony PCR. Plasmids were isolated with the E.Z.N.A.® Plasmid Mini Kit I (Omega Bio-tek, Norcross, GA, USA, Cat. No. D6943-03) and verified by Sanger sequencing (Beijing RuiBoxingke Biotechnology Co., Ltd., Beijing, China). All GS-system vectors were further validated by full-length nanopore sequencing (Beijing RuiBoxingke Biotechnology Co., Ltd., Beijing, China).

### *Construction of pY004, pY006, and pY009*

pY004 is the primary 4×myc-tagging entry clone. Its parent, PJG066, is a pGW7-derived binary vector in which the cauliflower mosaic virus (CaMV) *35S* promoter ($35S_{pro}$) has been replaced by the maize *ZmUBI* promoter ($UBI_{pro}$) and which carries a hygromycin resistance gene for eukaryotic selection (Yu et al., 2026). Because PJG066 contains an endogenous *Bsa*I site that would interfere with Golden Gate assembly, that site was removed by PCR amplification with primers PJG066-F/R to yield PJG097.

pY004 was then assembled from PCR fragments by seamless cloning: the gentamicin resistance gene *GentR* (GM-F/GM-R); the $UBI_{pro}$-4×myc driver (UBI-F/UBI-R); the lethal cassette (LC-F/R); and backbone fragments A (2.1 kb, BKA-F/R) and B (5.3 kb, BKB-F/R), all amplified from PJG097.

pY006 was generated by inserting *Paq*CI sites in inverted orientation adjacent to the *Bsa*I sites of *Bsa*I-linearized pY004 via seamless cloning, such that *Paq*CI digestion produces the same sticky ends as *Bsa*I (PCR with 4GS-F/R). To generate pY009, pY006 was linearized with *Sac*I and *Spe*I and assembled by seamless cloning with $35S_{pro}$ (amplified from pCAMBIA1301 with 35-F/R) and the 4×myc tag (amplified from pY004 with 4m-F/R), placing $35S_{pro}$ in place of $UBI_{pro}$.

### *pCAMBIA1301 backbone destination vectors*

The pCAMBIA1301-family destination vectors are based on pY002 (pCAMBIA1301_GSHCHC_K). The pCAMBIA1301 backbone was amplified from pCAMBIA1301-35S-MCS-35S-EGFP-Hyg with primers BKC-F/R. The *Hind*III-$35S_{pro}$-*Spe*I-3×HA-*Kpn*I module was produced by two rounds of PCR with primer pairs SHR1-F/R and SHR2-F/R, using pCAMBIA1301 as the initial template. The GS element (from pDONR207, GSC-F/R) was then combined with these fragments by one-step seamless cloning to yield pY002 and its derivatives.

pY003 and pY008 were produced by inserting $UBI_{pro}$ (amplified with U2H-F/R and U2-F/R, respectively) into pY002 linearized with *Hind*III and *Spe*I or *Hind*III and *Kpn*I, respectively. The N-terminal fluorescent-tag variants pY011, pY012, pY014, and pY015 were generated by recombining *Spe*I-*Kpn*I-linearized pY002 with a fluorescent tag: N-mNeonGreen (Shaner et al., 2013), mScarlet (Bindels et al., 2017), nYFP, or cYFP (Lee et al., 2008), amplified with 2NG-F/R, 2mS-F/R, 2nY-F/R, and 2cY-F/R, respectively.

The ethanol-inducible overexpression vector pY007 (pCAMBIA1301_GSHAHC_K) was built on the stable-transformation backbone pY003. The *alcR*–$35S_{Ter}$–$alcA_{pro}$–*mini35S* fragment (*alcR*, the

*Aspergillus nidulans* ethanol-responsive transcription factor; $35S_{Ter}$, the CaMV 35S terminator; $alcA_{pro}$, the ethanol-inducible *alcA* promoter; and *mini35S*, a minimal *35S* promoter driving the gene of interest) was amplified with EIO-F/R from pBBB-GRA-Cre/loxP (Zhao et al., 2011). pY003 was linearized with *Spe*I (a unique site immediately downstream of $UBI_{pro}$) and recombined with the fragment by seamless cloning, so that the inducible driver sits between $UBI_{pro}$ and the N-terminal 3×HA tag.

### *pGreenII-62sk backbone destination vectors*

For transient-expression studies in *N. benthamiana* or protoplasts, pGreenII-62sk-derived destination vectors were assembled by Golden Gate with *Paq*CI. pGreenII-62sk was linearized with *Spe*I and *Hind*III and recombined with the GS element and a fluorescent tag. The N-terminal tag vectors pY042–pY046 were built by amplifying the GS element from pY002 with 62NGS-F/R and pairing it with nYFP (62nY-F/R), cYFP (62cY-F/R), mTurquoise2 (62mT-F/R)(Goedhart et al., 2012), N-mNeonGreen (62NG-F/R), or mCherry (62mC-F/R). The C-terminal mNeonGreen vector pY048 was generated by amplifying both the GS element and mNeonGreen from pY045 with 62CGS-F/R and 62CNG-F/R, respectively.

### *Yeast two-hybrid destination vectors*

For yeast two-hybrid assays, pGADT7 and pGBKT7 were linearized with *Xho*I and *Bam*HI. The GS element was amplified with AGS-F/R (pGADT7) or BGS-F/R (pGBKT7) and introduced into the corresponding backbones by seamless cloning, yielding the prey vector pY061 and the bait vector pY062.

### *Bacterial-expression destination vectors*

Bacterial-expression destination vectors were built on pET and pGEX backbones. pET28a(+), pET30a, and pET49b were linearized with *Xho*I and *BamH*I and recombined with the GS element (amplified with 28GS-F/R, 30GS-F/R, and 49GS-F/R, respectively) by seamless cloning, giving pY071 (pET28a(+)-GS), pY072 (pET30a-GS), and pY074 (pET49b-GS). pGEX-6P-1, linearized with *Eco*RI and *Xho*I, was combined with the GS element (6PGS-F/R) in a *Paq*CI Golden Gate reaction to yield pY077.

To produce fluorescently tagged bacterial fusions, the sfGFP coding sequence (synthesized by Zhonghe Gene Co., Ltd., Tianjin, China, and cloned into pUC19) was amplified with sf-F/R. The GST-GS element was amplified from pY077 with CGG-F/R. A pColdIII-derived expression clone carrying a *Spe*I site inserted downstream of the TEE (translation-enhancing element) sequence was linearized with *Spe*I and *Xba*I and assembled with sfGFP and the two amplified fragments by seamless cloning; the resulting backbone pY081 carries an sfGFP-TEV site-GST-HRV 3C site-GS cassette.

The 6×His-HRV 3C-GS fragment was assembled from pY045 by two rounds of PCR with HGS1-F/HGS-R and HGS2-F/HGS-R. It was then recombined with *Hind*III-*Xba*I-digested pY081 in a *Paq*CI Golden Gate reaction to give pY082, in which the GST tag is replaced by a 6×His tag. Likewise, three rounds of PCR with SGS1-F/SGS-R, SGS2-F/SGS-R, and SGS3-F/SGS-R produced the S-tag-HRV 3C-GS fragment, which was inserted into pY081 to yield pY083.

Finally, pY071 was digested with *Xho*I and recombined with sfGFP-GST-GS (from pY081, GTG-F/R) or sfGFP-6×His-GS (from pY082, GTG-F/R) by seamless cloning. Sequence-verified clones were designated pY078 (pET28-GG-GS) and pY079 (pET28-GH-GS).

### ***Bidirectional promoter identification and characterization***

Bidirectional promoters were identified from *RiceSuperPIRdb* (Shang et al., 2023) under three criteria: head-to-head adjacent gene pairs on opposite strands, transcription start sites within 2 kb, and expression comparable to the rice reference gene *OsUbq* (*LOC_Os03g13170*).

The editability of the candidate intergenic region was predicted using CAPE, an open-source program available on GitHub (https://github.com/zhangtaolab/CAPE) (Zhou et al., 2023). Open chromatin data were obtained from the NCBI database (PRJNA705005); after quality control and low-quality read filtering, the high-quality reads were aligned to the rice reference genome IRGSP-1.0 using BWA (Li and Durbin, 2009). Histone post-translational modification (PTM) data were obtained from PlantDHS (Zhang et al., 2016). Precomputed sequence conservation scores were downloaded from PlantRegMap (Tian et al., 2019) and converted to BigWig with bedGraphToBigWig (Kent et al., 2010). Transcription factor binding sites (TFBSs) were predicted with FIMO v5.5.9 using the JASPAR 2024 CORE plants non-redundant position-frequency matrices (MEME format) (Rauluseviciute et al., 2024); sites with $p < 1.0 \times 10^{-4}$ were retained. CAPE then integrated open-chromatin accessibility, histone modifications, sequence conservation, and transcription factor binding sites to predict promoter editability.

The dual-fluorescent reporter vectors PFA467 and PFA491 were assembled by *Bsa*I Golden Gate cloning from pY002 linearized with *Hind*III and *Bam*HI. *35S*$_{pro}$*-mCherry-NOS*$_{Ter}$ (PA-F/R), *Spe*I-*mNeonGreen-35S*$_{Ter}$ (PB-F/R), *Spe*I-*mTurquoise2-35S*$_{Ter}$ (PD-F/R), and *pBD* (PC-F/R) were combined in one reaction; the resulting PFA467 constitutively expresses *mCherry* and drives *mNeonGreen* and *mTurquoise2* from the *pBD* promoter. PFA491 was derived by flanking *pBD* with a *mini35S* promoter on both sides through seamless cloning between *Spe*I-linearized PFA467 and *mini35S-pBD-mini35S* (MBM-F/R).

### *GSprimer primer-design skill*

GSprimer designs GS-system cloning primers from a gene identifier (default: the longest transcript) or a transcript ID, together with locally downloaded genome annotation and sequence files. For each input it anchors primers at the CDS, appends the GS adapter overhangs, verifies in-frame fusion with the 4×myc tag of GDVs, screens primer specificity genome-wide, and reports risks with concrete remedies.

#### CDS anchoring

The forward primer is positioned so that its 5' end coincides with the ATG, and the reverse primer so that its 3' end coincides with the last nucleotide of the stop codon, preserving the reading frame of the insert. Any permitted forward shift is a multiple of three nucleotides. A shift into the 5' UTR is allowed only when UTR sequence is available; a shift that starts inside the CDS (truncating N-terminal residues including Met) is heavily penalized and never reaches the top tiers. The strict ATG-stop pair is always retained as the reference candidate.

**Thermodynamic and sequence-composition criteria**

Each primer is screened for properties that govern amplification fidelity: (i) Length 19–24 nt and bulk GC content 40%–60%, with evenly distributed GC. Runs of >3 consecutive G or C are penalized. A 3' GC-clamp (≥ 3 G/C among the terminal 5 nt) is avoided because it promotes mispriming, whereas a single terminal G/C that closes the 3' end is mildly favored. (ii) Tm 55–60 °C with salt and $Mg^{2+}$ correction, and |ΔTm| between the pair < 2 degrees C. (iii) Secondary structure: self-dimer ΔG ≥ −6.0 $kcal \cdot mol^{-1}$ (≥ −5.0 at the 3' end), hairpin ΔG ≥ −4.0 $kcal \cdot mol^{-1}$, and cross-dimer between the two primers ΔG ≥ −6.0 $kcal \cdot mol^{-1}$. (iv) Low complexity is rejected: no mononucleotide runs > 4 nt and no dinucleotide repeats. The terminal 4–5 nt must be non-self-complementary and not complementary to the partner's 3' end, the main determinant of false priming. GSprimer imposes no amplicon-size window, because the CDS amplicon is set by the open reading frame.

**Tiering**

Each candidate pair is scored and assigned to Tier A, B, or C. Tier A requires the strict ATG-stop anchor (or a 3-nt-multiple UTR shift), LOW specificity risk, GC and Tm within range, ΔTm < 2 °C, no internal *Bsa*I/*Paq*CI site (or a synonymous wobble rescue), and an intact frame with no N-terminal truncation. Tier B tolerates minor deviations (a single borderline thermodynamic parameter, or a small but acceptable dimer or hairpin ΔG). Tier C is assigned to candidates with N-terminal truncation, HIGH specificity risk, or out-of-range Tm or GC that would compromise amplification or cloning. The ranked table lets the user pick a pair before adapter attachment.

#### Specificity screening and *Bsa*I/*Paq*CI scanning

Every candidate is checked in two layers. First, each primer is searched against the genome and transcriptome by BLAST (blastn-short), with a pure-Python seed-and-extend fallback when BLAST is unavailable. Second, an *in silico* PCR test verifies the pair: a primer is flagged as problematic only when its partner lies in the opposite orientation within an amplifiable distance, which sets the reported risk level. Perfect matches on different genes are rated HIGH; hits on isoforms of the same locus are expected and treated as LOW, because cloning all isoforms of the target is desirable. The amplicon is scanned on both strands for internal *Bsa*I/*Paq*CI sites; when found, GSprimer reports the exact codon positions and offers synonymous wobble edits that remove the site without changing the protein.

#### Template choice and PCR guidance

Genes with introns (multi-exon in the GFF3 annotation) are directed to a cDNA template, whereas intronless genes are directed to genomic DNA (no reverse transcription required). For the final ordered oligo, GSprimer confirms exactly two *Bsa*I sites (the two adapters). PCR recipe advice is auto-generated from fragment and primer traits: GC-rich templates call for a GC-enhanced high-fidelity polymerase with DMSO or betaine and touchdown PCR; long fragments call for a long-range polymerase and extended extension times; Tm-mismatched pairs are annealed at the lower Tm.

### *Cloning workflows of OsCaM61, OsMYL1, and OsbZIP83*

The end-to-end workflow was validated by cloning three rice genes: *OsCaM61* (LOC_Os01g59530), *OsMYL1* (LOC_Os01g50940), and *OsbZIP83* (*LOC_Os12g05680*). GSprimer was used to design GS-system-specific CDS-amplification primers; from the candidate pairs, the final primers were selected manually and validated. The CDS fragments of *OsCaM61* and *OsMYL1* were amplified by PCR, purified, and assembled with *Bsa*I-linearized pY004 by seamless cloning; the CDS of *OsbZIP83* was inserted into circular pY006 by Golden Gate using *Paq*CI. Sequence-verified entry and expression clones (EECs) were designated PFA454 (*OsCaM61*), PFA495 (*OsCaM61* without stop codon), PFA489 (*OsMYL1*), and PFA606 (*OsbZIP83*). These EECs were then assembled with destination vectors by the circular-to-circular Golden Gate integration strategy to produce the corresponding expression constructs, as detailed below. The CDS of *OsCaM61* was inserted into pGreenII-backbone destination vectors (pY044, pY045, pY046, and pY048) for subcellular localization in both rice protoplasts and *N. benthamiana* leaves. The CDS of *OsbZIP83* was inserted into pY061 and pY062 for Y2H, into pY014 and pY015 for BiFC in *N. benthamiana* leaves, and into pY042 and pY043 for BiFC in rice protoplasts. The CDS of *OsMYL1* was inserted into pY077 and pY081 for bacterial expression.

### *Transient expression in rice protoplasts*

Subcellular localization and BiFC assays were performed in rice protoplasts. Rice protoplasts were prepared and transformed by PEG/$Ca^{2+}$-mediated transfection as previously described (Chen et al., 2006; He et al., 2016). In brief, rice seeds were surface-sterilized and grown on half-strength MS medium in the dark at 26–28 °C for approximately 2 weeks. Protoplasts were isolated from the leaf sheaths of etiolated seedlings by enzymatic digestion with Cellulase RS (1.5% w/v; Yakult Honsha, Tokyo, Japan, Cat. No. MX7353) and Macerozyme R-10 (0.75% w/v; Yakult Honsha, Tokyo, Japan; Cat. No. MX7351). For each transformation, 10 µg of each purified plasmid was co-transfected into protoplasts, which were then incubated in the dark at 28 °C for 12 h before imaging.

### *Nicotiana benthamiana leaf agroinfiltration*

*N. benthamiana* seeds were germinated and grown in peat-based potting soil at 25 °C under a 14-h-light/10-h-dark photoperiod. DNA constructs were introduced into *Agrobacterium tumefaciens* GV3101 by the freeze–thaw method; pGreenII-derived constructs were transformed into the AGL1 strain carrying the *pSoup* helper plasmid. Transformants were selected on LB agar plates supplemented with rifampicin (25 µg/mL) and kanamycin (50 µg/mL). Three independent colonies per construct were verified by colony PCR, and one positive clone was grown for 16–20 h at 28 °C with shaking at 220 rpm in liquid LB containing the same antibiotics. Bacteria were harvested by centrifugation (6,000 rpm, 3 min), washed once with MMA buffer (10 mM MES, pH 5.6; 10 mM $MgCl_2$; 200 µM acetosyringone), and resuspended in 5 mL of freshly prepared MMA buffer. Suspensions were incubated with gentle shaking at room temperature for 1–2 h, and the $OD_{600}$ of each suspension was measured with a spectrophotometer. Bacterial suspensions, adjusted to an $OD_{600}$ of 0.2 with fresh infiltration buffer, were then manually infiltrated into the abaxial side of the third and fourth true leaves (counted acropetally, with the cotyledons designated leaves 1 and 2; three plants per infiltration) using a 1-mL syringe. Infiltrated plants were maintained in darkness for 24 h post-infiltration, followed by exposure to light (12–24 h). Leaf discs were harvested from the infiltrated area at 36–48 hours post-infiltration and subjected to confocal laser-scanning microscopy.

### *Confocal laser scanning microscopy*

Cells expressing fluorescently tagged fusion proteins were imaged on an LSM 980 confocal laser-scanning microscope (Carl Zeiss, Oberkochen, Germany). Fluorophores were excited and detected as follows: mTurquoise2, 405-nm excitation with emission collected at 455–526 nm; mNeonGreen, 488-nm excitation with 495–521 nm; YFP, 514-nm excitation with 525–566 nm; mScarlet, 543-nm excitation with 422–477 nm, 555–630 nm emission; mCherry, 543-nm excitation with 408–698 nm emission. When multiple fluorophores were imaged in the same sample, channels were acquired sequentially (multitrack mode) to minimize spectral bleed-through. Images were acquired and processed using ZEN software (Carl Zeiss).

### *Bacterial expression and purification of recombinant proteins*

The CDS of *OsMYL1* was cloned into the vector pY077-pGEX-6P-1_GS to generate GST–OsMYL1 and into the cold-shock vector pY081-pCold-GG_GS to generate sfGFP–GST–OsMYL1. The constructs were transformed into *E. coli* BL21(DE3), and transformants were grown in LB medium supplemented with carbenicillin (50 μg/mL; for pY077) or kanamycin (50 μg/mL; for pY081) at 37 °C until the $OD_{600}$ reached 0.6–0.8. Protein expression was induced with isopropyl β-D-1-thiogalactopyranoside (IPTG) under vector-specific regimes: for pY077 expression, IPTG was added to a final concentration of 0.5 mM and incubation continued at 16 °C overnight (~16–20 h); for cold-shock expression, cultures were pre-chilled at 15 °C for 1 h before induction with 20 μM IPTG, followed by incubation at 15 °C for 20–24 h.

For both constructs, cells were harvested by centrifugation at 8,000 rpm, 4 °C, for 10 min, and the pellet was resuspended in lysis buffer (50 mM Tris-HCl, pH 7.4; 400 mM NaCl; 10% v/v glycerol) containing phenylmethylsulfonyl fluoride (PMSF) at a final concentration of 1 mM. The suspension was lysed by high-pressure homogenization at 4 °C and 600 bar, and the lysate was clarified by centrifugation (8,000 rpm, 4 °C, for 1 h). The supernatant was loaded onto a gravity-flow column packed with Glutathione Beads 4FF (Smart-Lifesciences, Changzhou, Jiangsu, China; Cat. No. SA010025) pre-equilibrated with lysis buffer. The resin was washed with 4 column volumes of lysis buffer, and GST-tagged proteins were eluted with 7.5 mM reduced glutathione in lysis buffer.

Aliquots of uninduced culture, induced total lysate, soluble supernatant, and purification fractions (flow-through, wash, and elution) were mixed with 5× SDS-PAGE loading buffer and heated at 95 °C for 7 min. Proteins were resolved by SDS-PAGE using a 12–20% (w/v) linear gradient gel and visualized by Coomassie Brilliant Blue R-250 staining.

### *EMSA and dmEMSA assays*

The assays were performed as a single binding-and-gel protocol; the only differences between the two detection modalities were the probe/protein label and the gel-scanning program. Binding reactions (20 μL) contained the fusion protein (2 μg), a labeled probe (1 μM; 5'-end modified and annealed to the complementary strand), and EMSA buffer (10 mM Tris-HCl, pH 7.5; 50 mM KCl; 1 mM DTT; 5%

v/v glycerol; 5 mM $MgCl_2$ and 1 µg poly(dI·dC)). Reactions were incubated on ice for 25 min in the dark to avoid photobleaching. Loading dye without SDS was added before electrophoresis on a 4% (w/v) native PAGE gel in 0.5× TG buffer (25 mM Tris, 190 mM glycine); the gel was pre-run at 4 °C for 30 min, then run at 90 V for 65 min until the bromophenol blue reached the bottom, with the gel protected from light throughout. Specificity was controlled in both modalities using a 0.5–10× molar excess of unlabeled probe.

Two labeling and scanning configurations were used:

The probe carries a single label and the gel is imaged in a single channel in standard EMSA. In dmEMSA, the protein is expressed as an sfGFP fusion (sfGFP-OsMYL1) and the probe is 5'-Cy5-modified, so both partners are detected by in-gel fluorescence. The gel was imaged directly on a GE Amersham Typhoon 5. Channel 1 (protein, green) used a 488-nm laser with the sfGFP emission filter; Channel 2 (probe, red) used a 635-nm laser with the Cy5 670BP30 filter. Normal-scan mode with a pixel size of 100 µm minimized background. The two channels were false-colored green (sfGFP) and red (Cy5) and merged in ImageQuant TL; yellow indicates co-localization of protein and probe in the shifted complex. This dual-detection strategy follows the logic of shift-Western assays (Demczuk et al., 1993) but detects both partners by in-gel fluorescence of the tagged protein and the fluorescent probe, omitting the antibody step.
